# Wheat developmental traits as affected by the interaction between *Eps-7D* and temperature under contrasting photoperiods with insensitive *Ppd-D1* background

**DOI:** 10.1101/2020.09.10.290916

**Authors:** Priyanka A. Basavaraddi, Roxana Savin, Simon Griffiths, Gustavo A. Slafer

## Abstract

Earliness *per se* (Eps) genes are important to fine tune adaptation, and studying their probable pleiotropic effect on wheat yield traits is worthwhile. In addition, it has been shown that some Eps genes interact with temperature. We studied two NILs differing in the newly identified *Eps-7D* but carrying insensitive *Ppd*-D1 in the background under three temperature regimes (9, 15 and 18 °C) and two photoperiods (12 h and 24 h). *Eps-7D* affected time to anthesis as expected and the *Eps-7D*-*late* allele extended both the period before and after terminal spikelet. The interaction effect of *Eps-7D* × temperature was significant but not cross-over: the magnitude and level of significance of the difference between NILs with the *late* or *early* allele was affected by the growing temperature (i.e. difference was least at 18 °C and largest at 9 °C), and differences in temperature sensitivity was influenced by photoperiod. Rate of leaf initiation was faster in NIL with *Eps-7D*-*early* than with the *late* allele which compensated for the shorter duration of leaf initiation resulting in similar final leaf number between two NILs. *Eps-7D*-late consistently increased spike fertility through improving floret primordia survival as a consequence of extending the late reproductive phase.

## Introduction

Wheat development is critical for yield determination as it controls not only adaptation (i.e. the critical stage of anthesis must occur when conditions are best, minimising stresses during grain number determination and grain weight realisation; Fischer, 2011; Reynolds et al., 2012) but also the timing and rate of generation of structures that will become sources and sinks (González et al., 2005a; Whitechurch and Slafer, 2001). Indeed, wheat yield (as well as that of other grain crops) is the consequence of the balance between source- and sink-strength, in turn determined as the result of initiation, degeneration and rate of growth of leaves, tillers, spikelets, florets and grains. Genetic factors controlling the duration of the developmental phases would be expected to have pleiotropic effect on yield traits (Börner et al., 1993; Foulkes et al., 2004). Certainly, a number of studies have shown that modifying the duration of particular developmental phases either through genetic factors (Gawroński et al., 2014; Lewis et al., 2008; Ochagavía et al., 2018a; Pérez-Gianmarco et al., 2018; Prieto et al., 2018a) or environmental treatments (González et al., 2005a, 2003a, 2003b; Serrago et al., 2008; Steinfort et al., 2017; Wall and Cartwright, 1974) improves spike fertility; which in turn is a major determinant of wheat yield (Slafer et al., 2014; Würschum et al., 2018).

Time to anthesis in wheat encompasses various phases with different degrees of sensitivities towards cold temperature and daylength termed as vernalisation (*Vrn*) and photoperiod (*Ppd*) sensitivities, respectively. And the genetic factors responsible for such sensitivities are referred as *Vrn* and *Ppd* genes. The *Vrn*-sensitivity genes define the growth habit (*Vrn*-sensitive cultivars are winter wheats while *Vrn*-insensitive cultivars are spring wheats), while *Ppd*-sensitivity genes determine whether flowering will be earlier (cultivars with little or no sensitivity) or late (very sensitive cultivars) in spring. However, once the effects of *Vrn* and *Ppd* sensitivity genes are removed (because genotypes have insensitive alleles for all these genes or because plants are gown under long days after having been fully vernalised), genotypes may still exhibit differences in earliness of flowering. These genotypic differences are known as earliness *per se* (*Eps*) or intrinsic earliness (Slafer, 1996). Past wheat breeding has already ventured changing time to anthesis to expand adaptation and to maximise yield by positioning anthesis time to avoid yield penalties due to abiotic stresses (Araus et al., 2002; Richards, 1991). Then, major changes in anthesis time may not be as relevant as fine adjustments. The importance of Eps genes may be even higher than that of the major *Vrn* and *Ppd* sensitivity genes when the need is to fine adjust phenology because they normally have a relatively small effect (Bullrich et al., 2002; Griffiths et al., 2009; Lewis et al., 2008; Ochagavía et al., 2018b). Indeed, due to their relatively subtle effect, Eps genes may have gone undetected during the course of selection (Zikhali et al., 2014), and are mostly identified as QTLs (Zikhali et al., 2014). Although much lesser known, their possible pleotropic effect on yield components might be one of the reasons for their indirect selection (Alvarez et al., 2016).

Most of what is known of the identified Eps genes relates to their effects on time to anthesis. The importance of these genetic factors, like any other genes, to be used in breeding programmes is limited by the lack of understanding of their detailed effect on individual phases occurring before anthesis, and their possible influence on different yield attributes along the way. Although yield components are being determined during the whole growing season, some phases are more critical than others (Fischer, 2007; Slafer, 2003). Duration of phase before and after terminal spikelet (TS) may have completely different relevance for yield determination. Indeed, it is during the TS-anthesis phase that spike development controlling spike dry weight and spike fertility are determined (Abbate et al., 1997; Fischer, 2007; Halloran and Pennell, 1982; Serrago et al., 2008).

Some recent studies have shown the possible relevance of Eps genes not only in fine adjusting anthesis time, but also through affecting spikelet number (Alvarez et al., 2016) and grains per spike (Lewis et al., 2008). This is in line with the hypothesis that genes effecting developmental traits might alter the dynamics of organs initiated in response to changes in the duration (Ferrante et al., 2013; González et al., 2005b; Miralles and Richards, 2000; Prieto et al., 2018a, 2018b; Snape et al., 2001). The dynamics of organs such as tillers, spikelets and florets (resulting *a posteriori* in yield components) may well depend, at least in part, upon the time allocated for their development.

Despite Eps genes owe their name to the assumption that genotypic differences produced were “intrinsic” (*per se*) and therefore independent of the environment (Slafer, 1996), it was hypothesised to be temperature sensitive genes (Slafer and Rawson, 1995). The speculated Eps × temperature interaction (Appendino and Slafer, 2003; Bullrich et al., 2002; Lewis et al., 2008) was recently proven in few studies (e.g. Ochagavía et al., 2019; Prieto et al., 2020). However, what we collectively call Eps genes are consistent in their effect on time to anthesis, but could strongly differ in their effects on other traits. It could be possible that the temperature responses of each Eps be different in terms of type and magnitude of the response and this needs to be studied. Understanding whether temperature affects the functionality of each Eps is necessary to explore the kind of environment those Eps could be effective and beneficial.

Recently an Eps QTL on chromosome 7D was identified in wheat which was known to influence time to heading. Four NILs were generated from the cross Paragon (a modern UK commercial cultivar; e.g. (Wingen et al., 2017) and Baj (a CIMMYT cultivar, used frequently as check; e.g. (Mondal et al., 2016) both of which are spring type with no requirements of vernalisation. Paragon has the *Eps-7D*-*late* and *Ppd-D1b* alleles while Baj has the *Eps-7D*-*early* and *Ppd-D1a* alleles. Thus the four NILs comprised the four combinations of both alleles and had identical mixture of Paragon and Baj in the background. For simplicity of presentation of results, in the present paper we aimed to evaluate the direct effect of the *Eps-7D* alleles (comparing the performance of the NILs having always the *Ppd-D1a* allele) and the interaction with temperature at two contrasting photoperiods to quantify mainly the effect of *Eps-7D* on phenology as well as dynamics of organ development. The NILs were grown under three constant temperatures (9, 15 and 18 °C) and two very contrasting photoperiods (12 and 24 h). In a companion paper (Basavaraddi et al., submitted), we analysed to what degree the allelic form of the *Eps-7D* gene affect the sensitivity to photoperiod given by the strongest *Ppd* gene (*Ppd-D1*) and its interaction with temperature as well as whether the allelic form of *Ppd-D1* in the background modifies the effect of *Eps-7D* and its interaction with temperature.

## Results

Time to anthesis was inversely related to both growing temperature (longest at 9 °C and shortest at 18 °C) and photoperiod (longest at 12 h and shortest at 24 h) (Fig. 1), the latter even though all lines carry the insensitive photoperiod allele in chromosome 1D (*Ppd-D1a*). Although these two direct effects of temperature and photoperiod are expected we also found a significant interaction between them (Fig. 1c), that was not simply a reflection of the temperature effect on development as the difference between short and long photoperiod was largest in the intermediate temperature. This interaction reflects the fact that sensitivity to temperature was stronger under long than under short photoperiod (*cf*. Fig. 1a and 1 b). The interaction was significant but not cross-over: the NIL with the *Eps-7D*-*late* allele was always later to flower than that with the *early* allele (Fig. 1), but the magnitude and level of significance of the difference between NILs with the *late* or *early* allele was affected by the growing temperature (i.e. difference was least, and non-significant under SD, at 18°C and largest and clearly significant at 9°C; Fig. 1a, b). The effect of the *Eps-7D* gene did not show any interaction with photoperiod (Fig. 1c) and therefore the magnitude of difference between *Eps-7D*-*late* and *early* NILs were similar at both photoperiods, but when considered within each particular environment, the differences were more significant under long than under short days (Fig. 1a, b).

**Figure 1.**
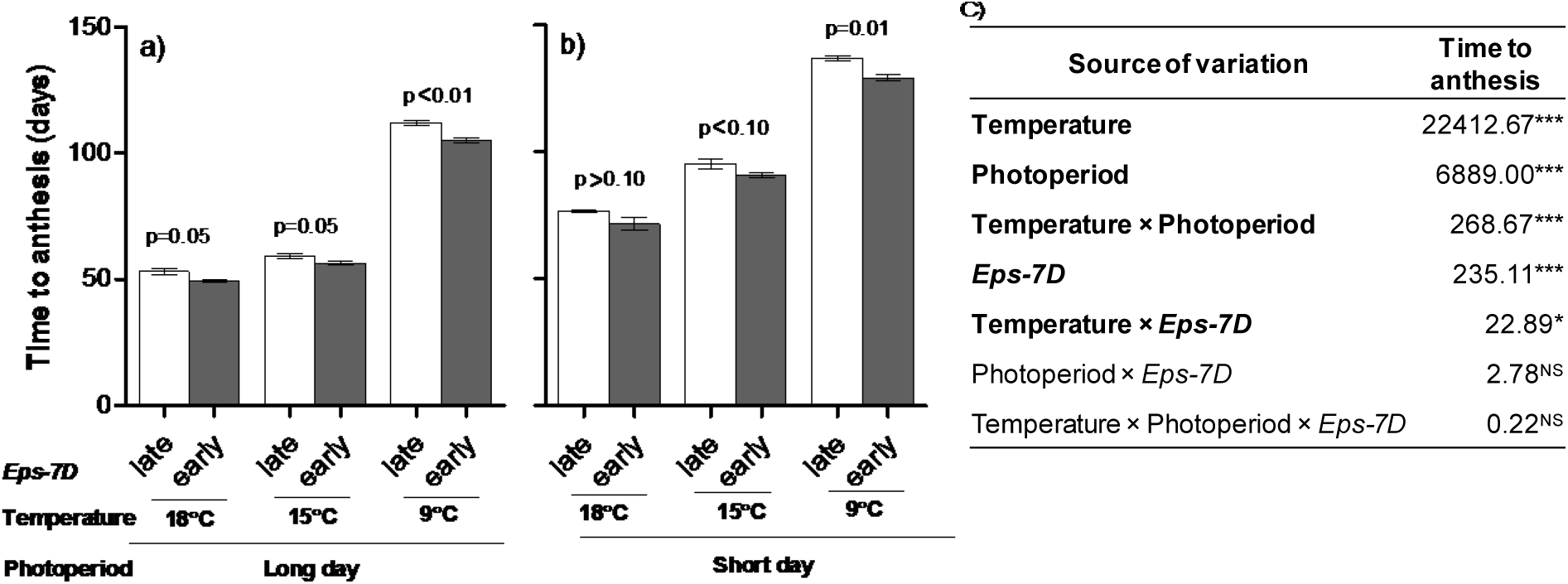
Duration of whole phase from seedling emergence to anthesis for the lines carrying *Eps-7D*-*late* (open bars) or -*early* (closed bars) on *Ppd*-D1a background under three growing temperatures at long day (a) and short days (b). Error bars indicate the SEMs of the mean and the “P” values stand for the level of significance exclusively due to the action of the *Eps-7D* gene within each temperature and photoperiod condition. The output (mean squares) of the three-way ANOVA for time to anthesis (days) is included on the right (c). Significance level * p < 0.05; ***p < 0.001; NS= non-significant.

The effects of temperature and photoperiod on time to anthesis were also seen for the two component phases considered here: both time from seedling emergence to TS (when all leaves and spikelets are initiated) and from then to anthesis (i.e. the late reproductive phase of stem elongation, LRP) were longer under low temperatures and short photoperiod than under warm temperatures and long photoperiod (Fig. 2). However, (i) even though both phases were clearly sensitive to the growing temperature, their sensitivity was not the same: duration from seedling emergence to TS responded to temperature less markedly than duration of the LRP (cf. differences between Fig.2a and b with Fig.2c and d, taking into account the different scales); and (ii) alike for the whole period to anthesis the sensitivity to temperature was stronger under long than under short days for both phases (Fig. 2). Regarding the specific effect of the *Eps-7D* gene, the NIL with the *Eps-7D*-*late* allele tended to have longer phases both from seedling emergence to TS and from then to anthesis across all growing conditions (Fig. 2).

**Figure 2.**
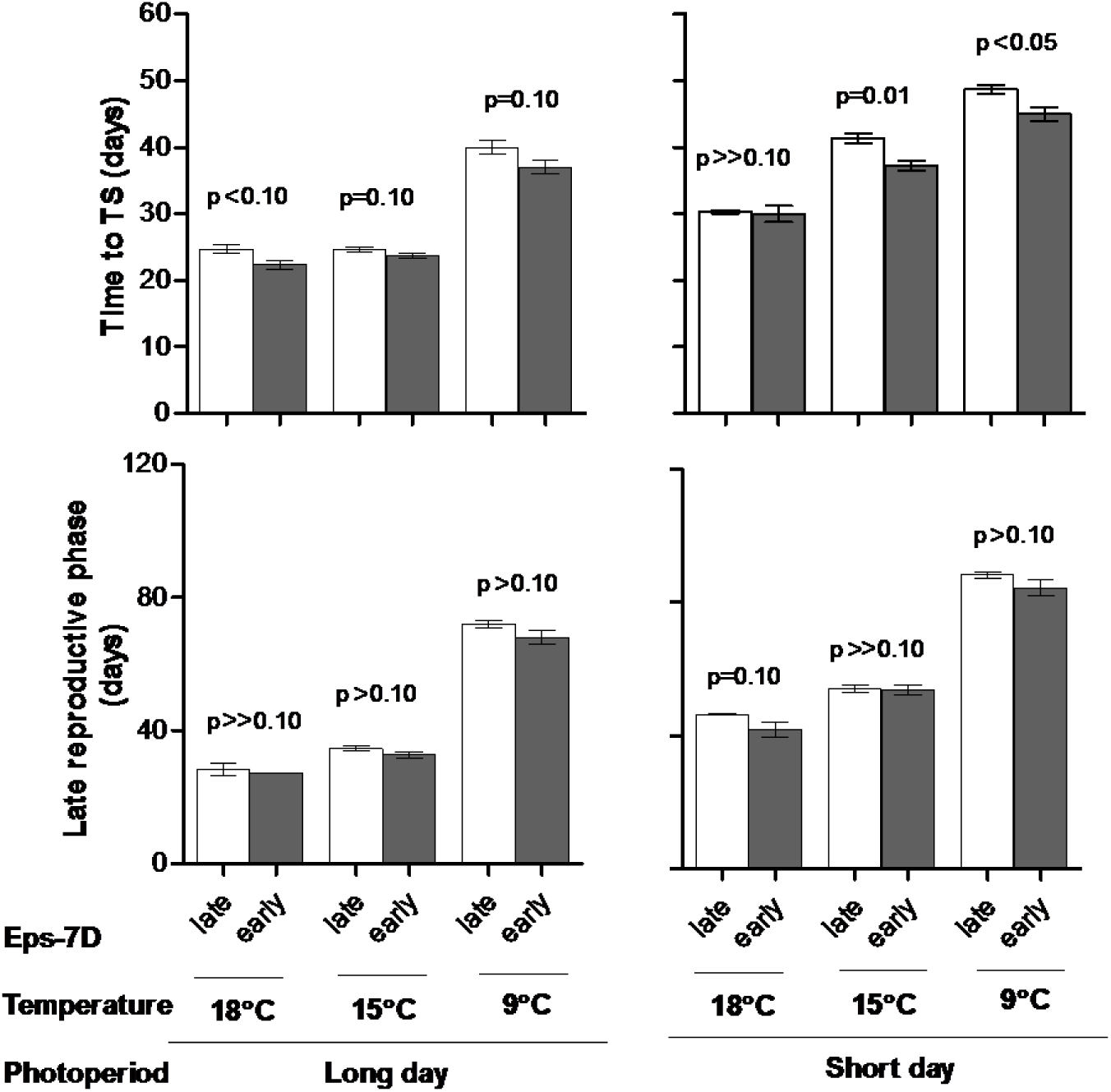
Duration of phase from seedling emergence to TS (upper panels) and time from then to anthesis, late reproductive phase (lower panels) for the lines carrying *Eps-7D*-*late* (open bars) or *early* (closed bar) on *Ppd*-D1a background under long (left panles) and short day (right panels) at three temperatures. Error bars indicate the SEs of the mean and the “P” values stand for the level of significance exclusively due to the action of the *Eps-7D* gene within each temperature and photoperiod condition.

However, as the effect on the whole period from seedling emergence to anthesis was subtle, that on the duration of each of its component phases was naturally even smaller and most differences became non-significant with the two-way ANOVA analyses done for each growing condition; particularly for the LRP (Fig. 2). But looking at the relationship between the duration of the total time to anthesis and its component phases it seems clear that both were at least equally important, not only reflecting the differences between growing conditions but also the effects of the *Eps-7D* gene (Supplementary Fig. S1). Thus, even though most differences between NILs with *Eps-7D*-*early* and -*late* alleles were non-significant for the LRP (Fig. 2c, d), it can be seen that the magnitude of the shortening of the phases produced by the effect of having the *Eps-7D*-*early* allele was similar in relative terms for both phases (averaging across the six growing conditions the duration of the phase to TS and that of the LRP was 2.5 and 3 d earlier, respectively in the NIL with the *Eps-7D*-*early* than with the -*late* allele).

Final leaf number was not significantly affected by temperature or the *Eps-7D* gene (Table 1). Thus, any effects of these two factors on the duration of the vegetative phase of leaf initiation (virtually from sowing to seedling emergence or soon after it; see below) would have been compensated by opposite effects on the rate of leaf initiation. Photoperiod effect on FLN was small but clear; averaging across temperatures and *Eps-7D* alleles plants developed slightly less than 1 additional leaf if grown under short photoperiod. This means that when plants were exposed to long days they immediately reached floral initiation at seedling emergence (as there would be 4 leaf primordia in the embryo and a couple would have been initiated between sowing and seedling emergence) whilst at short days it took an additional plastochron to reach floral initiation, a difference that was very slight as expected (all plants were insensitive to photoperiod regarding the major gene *Ppd*-D1).

**Table 1.** Effects of the *Eps-7D* gene on final leaf number (FLN), rate of leaf appearance (RLA; estimated as the slope of the linear regression of leaf number vs thermal time), and the coefficient of determination for that regression (r^2^), when grown under two contrasting photoperiods (12 and 24 h) and three temperatures

| Growing conditions | | Allele at *Eps-7D* | FLN | RLA (leaves d<sup>-1</sup>) | $r^2$ |
| --- | --- | --- | --- | --- | --- |
| Long day | 18 °C | Late | 6.2 ± 0.1 | 0.142 ± 0.003 | 0.953*** |
|  |  | Early | 6.0 ± 0.0 | 0.149 ± 0.005 | 0.923*** |
|  | 15 °C | Late | 6.0 ± 0.0 | 0.122 ± 0.001 | 0.986*** |
|  |  | Early | 6.0 ± 0.0 | 0.131 ± 0.001 | 0.983*** |
|  | 9 °C | Late | 6.0 ± 0.0 | 0.083 ± 0.001 | 0.980*** |
|  |  | Early | 6.0 ± 0.0 | 0.083 ± 0.001 | 0.968*** |
| Short day | 18 °C | Late | 7.0 ± 0.0 | 0.126 ± 0.001 | 0.983*** |
|  |  | Early | 6.6 ± 0.1 | 0.130 ± 0.002 | 0.975*** |
|  | 15 °C | Late | 7.0 ± 0.0 | 0.083 ± 0.001 | 0.985*** |
|  |  | Early | 6.9 ± 0.1 | 0.087 ± 0.001 | 0.984*** |
|  | 9 °C | Late | 6.7 ± 0.2 | 0.066 ± 0.001 | 0.959*** |
|  |  | Early | 6.1 ± 0.1 | 0.072 ± 0.001 | 0.977*** |

\*\*\*All linear regressions of leaf number vs time were highly significant ( $P < 0.001$ ; $n = 10-25$ , depending on the temperatures and photoperiod as leaf number was determined thrice a week)

The initiated leaves always appeared at a reasonably constant pace (as indicated by the very high coefficients of determination of the linear relationship between leaf number and time; r^2^>0.92, n≥10; Table 1). The rate of appearance of these leaves was positively affected by temperature and photoperiod (the higher the temperature or longer the day the faster the rate of leaf appearance; Table 1). The *Eps-7D* gene also affected slightly but consistently the rate of leaf appearance, appearing faster in NIL with the *Eps-7D*-*early* allele than the one with *late* allele, with the exception of plants under long days and 9 °C in which the rates of leaf appearance of the NILs did not differ (Table 1).

As floral initiation occurred at seedling emergence or just 1 plastochron later (see above), we could only collect data revealing the dynamics of spikelet initiation (and estimate from that dynamics the spikelet plastochron). Spikelets were initiated at a more or less constant rate whose actual value was rather similar (and few differences were not consistent) for NILs with the *early* or *late* allele in *Eps-7D*, and in all cases clearly slower at 9 than at 15 or 18 °C and slower under short than under long days (Fig. 3).

**Figure 3.**
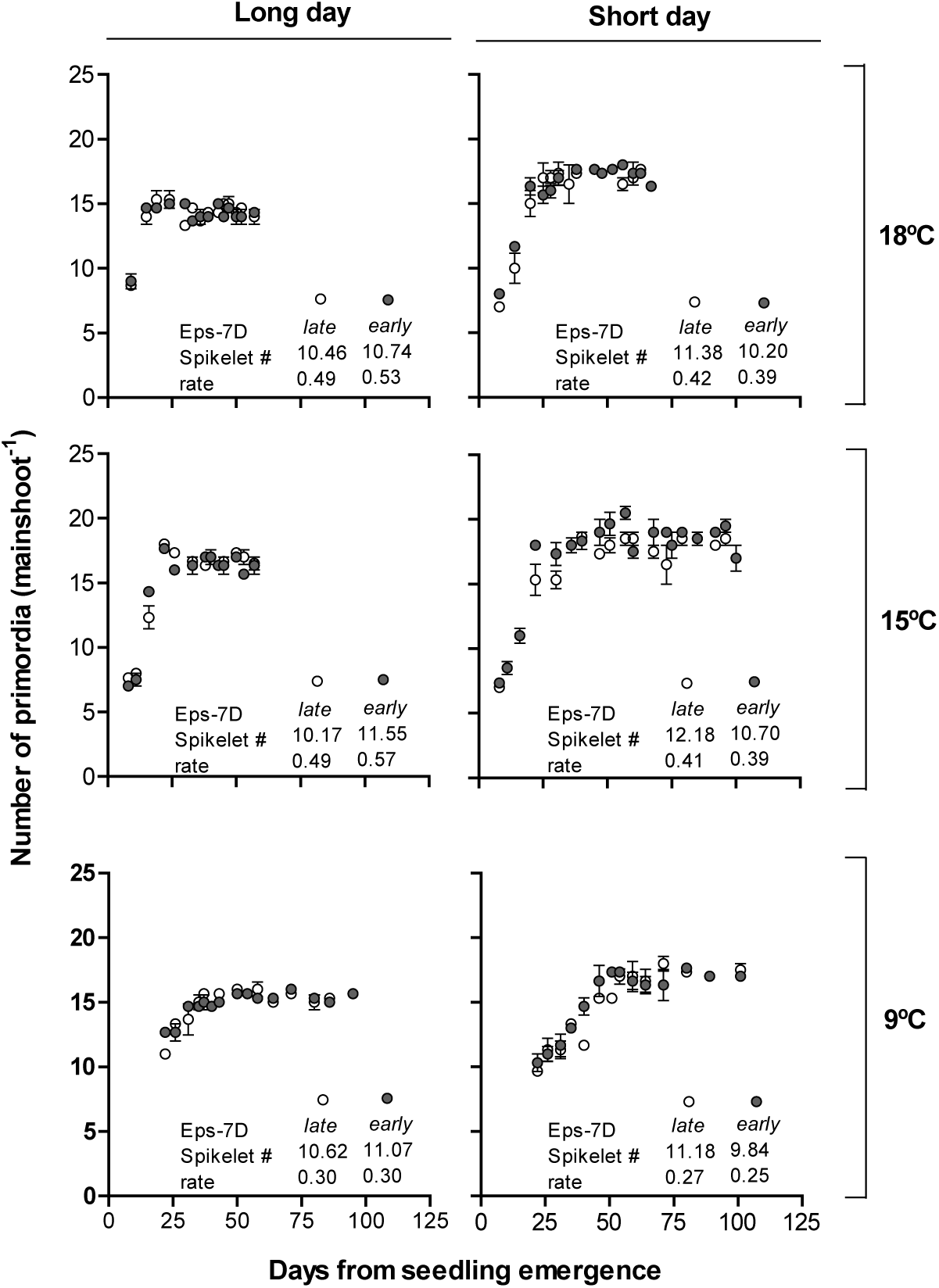
Relationship between number of primordia and days from seedling emergence for *Eps-7D*-late (open circles) and early (closed circles) under long (left panels) and short days (right panels) at 18 (upper top panels), 15 (middle panels) and 9 °C (bottom panels). Inside each panel are the total number of spikelet primordia and rate of spikelet initiation (spikelet primordia per day).

The dynamics of floret development was recorded for all the initiated florets within apical, central and basal spikelets that reached a developmental stage of W4.5 until they either reached W10 (fertile floret) or die. Floret 1 (most proximal floret to rachis) in both *Eps-7D*-*late* and *early* lines reached the stage of fertile floret (W10) under all three temperatures and two photoperiods, while F4 (the most distal floret consistently reaching at least the stage W4.5) has never reached to a stage close to W10 in any of the growing conditions (Supplementary Fig. S2). Then to understand the effects of treatments on spike fertility, we concentrated the results on the fate of the second and third florets from the rachis (F2 and F3 respectively) which were those responsible for the differences in number of fertile florets per spike at anthesis. Alike what was described for the initiation of spikelets, the rates of floret development were affected by the growing conditions. Florets developed much faster at 18 than at 9 °C but also the opposite was true with the duration of the period of floret development: shortest and longest at 18 and 9 °C, respectively (Figs. 4, S2). Photoperiod did not affect noticeably the rate of floret development but did modify the duration of the period of floret development (Figs. 4, S2).

**Figure 4.**
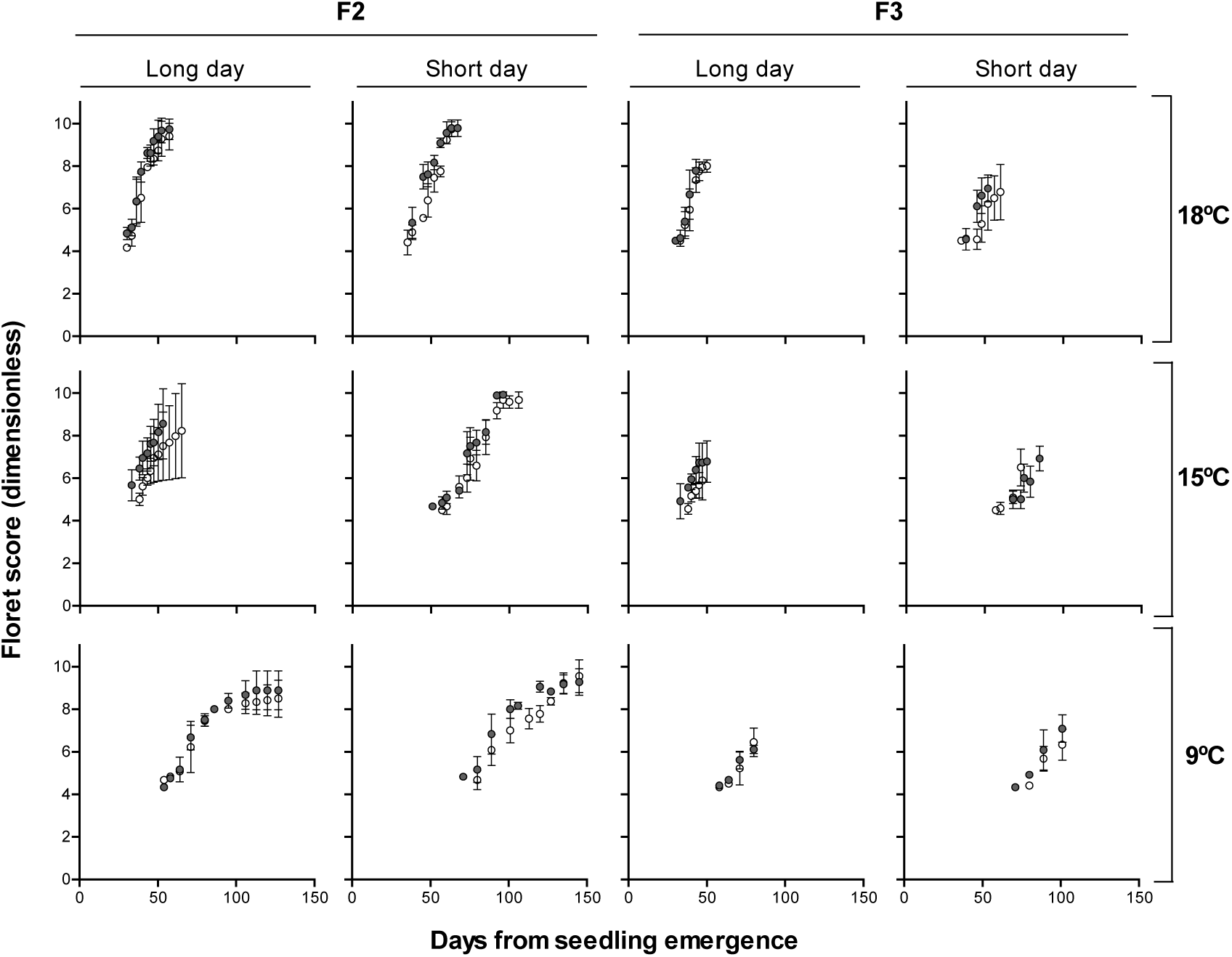
Relationship between floret development (floret score of the Waddington scale proposed by Waddington et al. 1983) and days from seedling emergence for *Eps-7D*-late (open circles) and early (closed circles) for floret F2 (left panels) and F3 (right panels) under long and shot day at 18 (upper panels), 15 (middle panels) and 9 °C (bottom panels). The error bars are SEs of means of floret scores from apical, central and basal spikelets.

**Figure 5.**
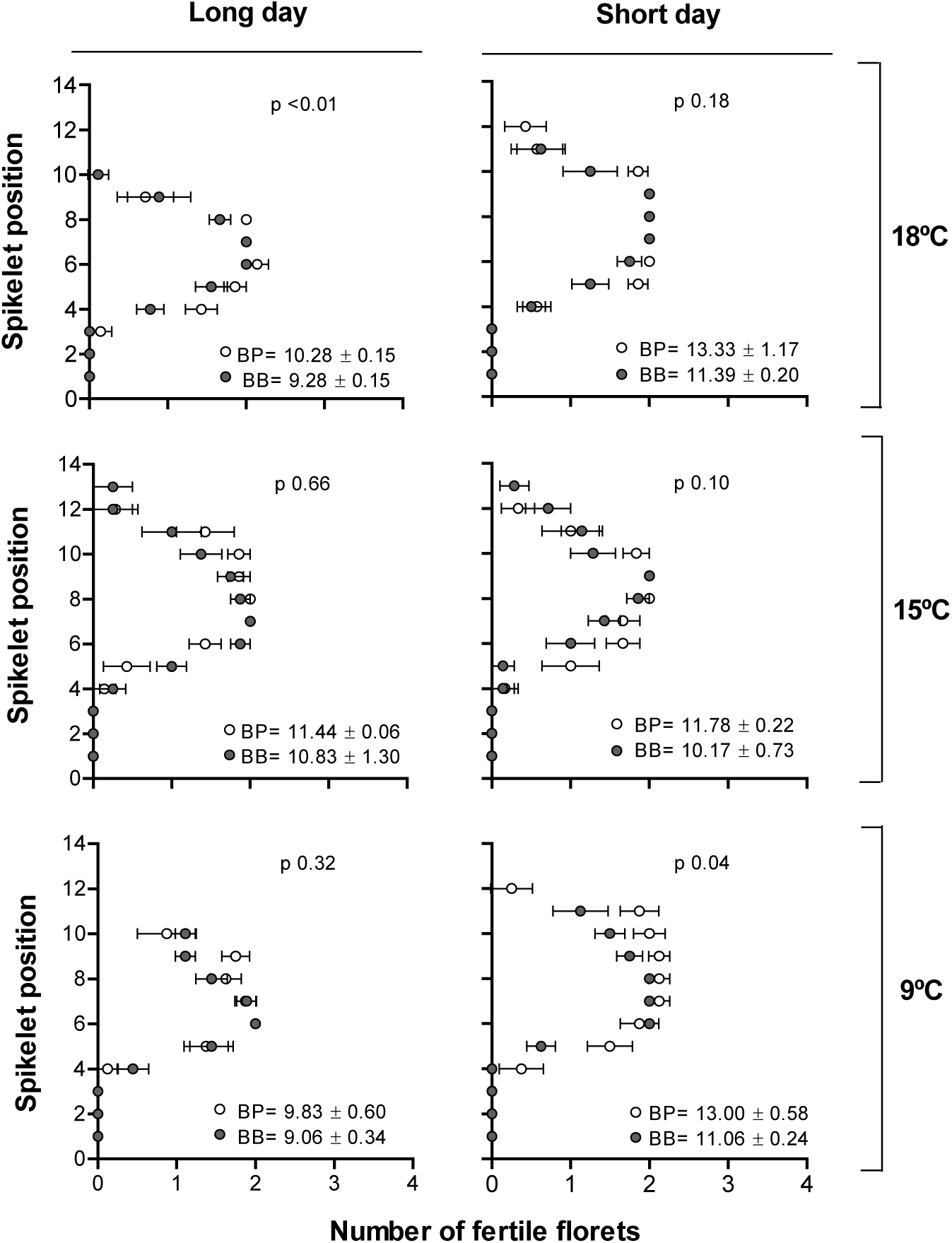
Number of fertile florets at anthesis per spikelet from basal to terminal spikelet for *Eps-7D*-*late* (open circles) and -*early* (closed circles) NILs under long (left panels) and shot days (right panels) at 18 (upper panels), 15 (middle panels) and 9 °C (bottom panels). Inside each panel are the fertile florets per spike ± SEs and p value.

Regarding the effect of the *Eps-7D* gene, Floret 2 was initiated more or less at the same time for both *Eps-7D*-*late* and -*early* under long day in all the three temperatures but under short day *Eps-7D*-*early* tended to initiate the F2 earlier and had faster development compared to *late* allele (Fig. 4). Under long day F2 reached W10 at 18°C for both *Eps-7D*-*late* and -*early* alleles, while one third of the florets F2 in *Eps-7D*-*late* reached W10 under lower temperatures (15 and 9 °C) and F2 from *Eps-7D*-*early* aborted when they had reached the W8.5 stage (green anthers). None of the F3 florets reached W10 regardless of whether the lines had the *Eps-7D*-*late* or -*early* alleles and therefore the effect of the *Eps-7D* gene was inappreciable. Even though the F4 florets did never reach the stage of fertile florets they attained higher floret score when the line had the *Eps-7D*-*late* allele, especially under short day conditions (Supplementary Fig. 2).

Spike fertility was not consistently affected by temperature (because of the opposite effects of this factor in the rate and duration of floret development, see above); and was higher in short than in long days by virtue of the photoperiod effect on duration of floret development (Fig. 6). The *Eps-7D* gene had an effect on the number of fertile florets per spike as the NIL with the *late* allele showed a consistent trend (though not always statistically significant) to have more fertile florets than the NIL with the *early* allele (Fig. 6).

The overall direct effect of *Eps-7D* gene on the number of fertile florets was much higher than the direct effect of temperature and *Eps-7D* × temperature interaction effect (F ratio was 8.50, 5.61 and 0.65 for *Eps-7D*, temperature and their interaction respectively). In that the averaging across the temperature the *Eps-7D*-*late* had almost c. 1 extra fertile floret per spike than that of early allele under LD and the difference doubled under short photoperiod. The huge effect of temperature on the phenology was not reflected in the fertile floret as temperature also affected the rate of floret development (similar to rate of leaf appearance and spikelet primordia initiation explained above) meaning longer duration of floret development due to low temperature did not allow more florets to advance towards fertile stage rather development of each floret was significantly slow (e.g. F1 took 22 d and 74 d at 18 and 9 °C, respectively under LD to advance from W4.5 to W10 for *Eps-7D*-*late* allele).

## Discussion

Although the main focus of this study was on the effects of this newly reported *Eps-7D* gene on developmental processes and whether or not those effects were affected by the growing temperature, we also reported the effects of temperature, photoperiod and their interaction on these developmental processes. As the temperature × photoperiod and *Eps-7D* × temperature interactions were significant (but that of *Eps-7D* × photoperiod and the triple interactions were not), we firstly discussed briefly the effects of the environmental factors and then those of the Eps and its interaction with temperature.

### Temperature, photoperiod and their interaction

In general, developmental rates were faster (reducing the length of both the whole cycle to anthesis and its component phases occurring before and after terminal spikelet) under high than under low temperature conditions. This overall effect is in line with the recognised universal effect of temperature on accelerating developmental processes not only in wheat (Slafer and Rawson, 1994a; John and Megan, 1999); as well as in and other crops (Parent and Tardieu, 2012) and other unrelated organisms (Gillooly et al., 2002). Also the rate of leaf appearance (that was constant for all leaves, as expected when FLN is less than 8; (Ochagavía et al., 2017; Slafer and Rawson, 1997) was positively responsive to temperature; as has been known for a long time (e.g. Miglietta, 1989; Slafer and Rawson, 1997). As temperature accelerated the rate of primordia initiation we found a sort of compensation with the acceleration of development (i.e. phases are shorter but primordia are initiated faster under higher temperatures). Consequently, not clear effects of temperature were evident for the final leaf number, the number of spikelets per spike or the number of fertile florets per spike, again as expected from this universal effect of temperature on rates of phenological development and of initiation of primordia during the corresponding phenological phases (Slafer and Rawson, 1994a).

There was a direct effect of photoperiod on time to anthesis, that was not restricted to the phase from seedling emergence to TS as the LRP was also affected by the exposure to contrasting day lengths (in line with previous evidences in the literature showing that the LRP can be highly sensitive to photoperiod; González et al., 2005b, 2003; Pérez-Gianmarco et al., 2018). As NILs had the insensitive allele for *Ppd*-D1 gene (*Ppd*-D1a), which is the insensitivity gene frequently reported to have the strongest effect (e.g. Langer et al., 2014; Pérez-Gianmarco et al., 2018), we did not expect large differences between growing the plants at short or long photoperiod. However, the NILs would have sensitive alleles in the *Ppd*-1 loci on A and/or B genome. These genes produce responses that are frequently less noticeable than *Ppd*-D1, but still significant (Bentley et al., 2011; Pérez-Gianmarco et al., 2018; Shaw et al., 2013, 2012). Again as expected from the literature, photoperiod effects on the rate of phenological development is not paralleled by concomitant effects on the rate of leaf initiation and therefore the final number of leaves was increased under short days (Slafer and Rawson 1994b). Long photoperiod not only reduced FLN but also accelerated the rate of leaf appearance (Mosaad et al., 1995; Slafer and Rawson, 1997) both factors contributing to the shortening of the time to anthesis produced by the extended photoperiod.

Beyond the direct effects of temperature and photoperiod discussed above, in the present study there was a clear temperature x photoperiod interaction. For instance, analysing in detail the responses to temperature in the contrasting photoperiods there were particularities that are worth noticing. The length of the phase under long day were similar for 15 and 18 °C while it differed clearly under short day between these temperatures showing shorter phase at 18 than at 15 °C indicating that the probable T_optimum_ for development under long days is lower than that under short day. This was all the more so when looking at the time to TS but not so much when LRP was considered, which is in line with the fact that cardinal temperatures would increase with the stage of development (Rahman and Wilson, 1978; Slafer and Savin, 1991; Slafer and Rawson, 1995). The fact that photoperiod affect the temperature response has been described several times not only for wheat (Kiss et al., 2017; Slafer and Rawson, 1996) but also for barley (Hemming et al., 2012; Karsai et al., 2013).

### *Eps-7D* and *Eps-7D* × temperature interaction

In line with the previous knowledge about other known Eps genes, the *Eps-7D* studied here also had subtle through consistent and significant effects on time to anthesis (Ochagavía et al., 2018b, 2019; Zikhali et al., 2014). This is not surprising as even though each Eps gene would have different mechanisms of action, by definition they all result in relatively small differences in time to anthesis or heading (Griffiths et al., 2009; Zikhali et al., 2014) to the degree that many times may be undetectable if photoperiod and vernalisation requirements are not fully satisfied (Zikhali et al., 2014). There are very fewer studies on detailed effect of Eps genes on pre-anthesis and, unlike with the overall time to anthesis, they vary in their conclusion on whether Eps affect early or late stages of development. While the study by Lewis et al. (2008) reported that the effect of Eps-A^m^l on time to anthesis was mainly due to its effect on the duration of early developmental phases until terminal spikelet, others (Ochagavía et al., 2018) reported varying effect of Eps-D1 on all the three phases, vegetative, early reproductive and late reproductive, depending on the cross (genetic background). The *Eps-7D* we characterised in the present study (with *Ppd*-D1a in the genetic background) was found to affect the duration of both the early phase from seedling emergence to TS as well as that of the LRP, similarly to what was reported for the Eps-D1 by Ochagavía et al. (2018). The effect of *Eps-7D* on time to anthesis was related to both number and rate of leaf appeared in that the NIL with *Eps-7D-late* allele had slightly more leaves developed that appeared slightly slower.

Considering that the NILs had similar FLN might seem like effect of *Eps-7D* on phenology was realised much later during the development (after flag leaf initiation). Indeed, the dissection of apex stipulated that the *Eps-7D* affected development since early reproductive phase. The rate of leaf appearance was affected by *Eps-7D* allele which resulted in *Eps-7D*-*early* allele to have similar FLN as that of *late* allele for a shorter duration. This implies a different mechanism regarding leaf development than what was shown for the Eps-D1; which affected time to anthesis through mainly affecting time from flag leaf emergence to anthesis (Prieto et al., 2020).

Improvements in spike fertility may be possible with either lengthening the LRP (with no compensation from the change in the rate of development, so that more florets may become fertile) and/or increasing spike dry weight at anthesis (which could be in turn the result of lengthened LRP or increased dry matter partitioning; Slafer et al., 2015). Changes in spike dry weight are uncertain with minor differences in phenology (unless partitioning was altered) and differences in spike fertility would be very subtle which would mainly be the result of the efficiency (Prieto et al., 2020 and references quoted there in). The consistent trend observed in the present study for the *Eps-7D*-*late* allele to produce more fertile florets per spike than the *early* allele was result of couple of extra florets in the distal position (F2 and F3 in this case) that continued developing for a slightly longer time as a consequence of the slightly lengthened LRP. Effect of *Eps-7D* on the duration of floret development did not alter number of florets primordia produced but altered floret survival which is strongly supported by other studies where major or minor differences in length of floret development phase resulting in differences in spike fertility was not through number of floret primordia produced (Prieto et al., 2020 and refernecs quoted there in). There was huge difference in duration of floret development between 18 and 9 °C but this did not generate similar improvement in fertile florets per spike at the low temperature because the driving force for decelerating the rate of development during the LRP was also decelerating the rate of floret development.

Further, in the present study there was clear interaction effect of *Eps-7D* × temperature on the phenology. Although temperature accelerates development of all phases in all crops (see above) that only means that there would be no cases of insensitivity, but genotypic variation in sensitivity has been shown since long time ago (Atkinson and Porter, 1996; Rawson and Richards, 1993; Slafer and Rawson, 1995). At least in part, the genotypic variation in sensitivity to temperature might reflect the interaction of Eps genes with temperature (Slafer, 1996). The interaction we found in this study between *Eps-7D* and temperature was not as obvious as to observe the inverse ranking of *Eps-7D*-*late* and *early* allele at varying temperature, but clear differences in the magnitude of the effect of the *Eps-7D* allele at different temperature. To the best of our knowledge such interaction had been only recently shown in hexaploid wheat for the Eps-D1 (Ochagavía et al., 2019), although it had been recognised time ago in diploid wheat (Bullrich et al., 2002), and now we expand the concept within commercial wheat germplasm to the new *Eps-7D*. Both the NILs carrying either *Eps-7D*-*late* and *early* accelerated the rate of development when the temperature was increased but the *Eps-7D*-*early* had higher sensitivity to temperature than the *late* allele which made *early* allele to have much shorted phenology than the *late* allele. Alleles of Eps genes might have different optimum temperatures which shows differences in earliness by *early* or lateness by *late* allele under various temperatures (Appendino and Slafer, 2003).

## Materials and methods

The experiments were conducted under controlled conditions in growth chambers (GER-1400 ESP, Radiber SA, Spain) at the University of Lleida, Spain. The pots (200 cm^3^) were filled with approximately 120-125 g of mixture of 70% soil and 30% peat. Two seeds were sown in each pot at uniform depth and were kept under dark at room temperature until seedling emergence. And only one seedling was retained per pot before shifting the pots to the growth chamber. Extra pots were sown to select 54 pots per NIL for each chamber which had uniform seedling emergence to avoid differences in plant development before the start of the experiment. Pots were watered once or twice a week based on the growth stage/water requirements/treatment. Micro and macro nutrients were provided through irrigation at 4-leaf stage in all growing conditions. Pots were rotated once a week within each chamber throughout the experimental period to eliminate any spatial variation causing differences in micro-environment within the chambers.

Treatments consisted of a factorial combination of four near isogenic lines (NILs) differing in the alleles of both *Eps-7D* (*Eps-7D*-*early* and-*late*) and *Ppd-D1* (*Ppd-D1a* and *Ppd-D1b*); two photoperiod conditions and three temperatures regimes. The NILs were derived from the cross Paragon and Baj carrying either *Eps-7D*-*late* and *Ppd-D1b* from Paragon or *Eps-7D*-*early* and *Ppd-D1a* from Baj. In this paper we focused on the effects of the *Eps-7D* gene and all NILs had the insensitive allele for this major *Ppd* gene (*Ppd-D1a*), and in the companion paper, we explored whether the sensitivity to photoperiod may affect the Eps-D7 (and Eps-D7 x temperature) effects. The plants were grown under either 12 or 24 h photoperiod (short day, SD and long day, LD, respectively), the treatment of LD having only half of the lights on so that daily radiation was the same for both photoperiod conditions. Three constant temperature regimes (9, 15 and 18 °C) were imposed under each of the two photoperiods from seedling emergence to anthesis. Nine randomly chosen plants per NIL in each of the six temperature × photoperiod conditions were marked at one leaf stage to record the dynamics of leaf appearance until the flag leaf was fully emerged. These plants were arranged in a completely randomise design with 9 replicates (each replicate being an individual plant). The leaf appearance was recorded three times a week for plants under LD and at least twice a week for plants under SD at all the temperatures following the scale proposed by Haun et al. (1973). The same plants were used to map the fertile florets (number of fertile florets at each spikelet) per spike at anthesis where florets that had either hanging anthers or were at least at the green anther stage were considered to be fertile. On all plants we measured (i) the phenological stages of flag leaf emergence (DC39), heading (DC59) and anthesis (DC65) by visual observation following the scale of Zadoks et al. (1974). The dates for each stage were recorded after observing the stage in number of representative plants in each NIL. The rest of the unmarked plants (45 in each combination of NIL x photoperiod x temperature) were also arranged in a completely randomised design and were sampled at regular intervals (depending on temperature and photoperiod treatment) to dissect and record the apex stages and number of primordia until the stage of terminal spikelet (TS), and from then to anthesis dissecting particular spikelets to determine the number and stages of each floret primordia. Three plants (replicates) per NIL within each treatment were sampled every time. Number of spikelet primordia was calculated *a posteriori* by subtracting final leaf number from the total number of (leaf and spikelet) primordia recorded until TS. For the determination of stages of development of the spike and florets we used the scale proposed by Waddington et al. (1983).

Nine plants per NIL that were reserved for recording the leaf appearance were sampled at anthesis, where the final number of fertile florets in each spikelet of the main shoot spike was determined. The florets were numbered F1 to Fn based on their position with respect to rachis, F1 being the most proximal to, and Fn the most distal from, the rachis. Wheat displays asynchronous development of florets across different spikelets of the spike, so dissection was carried out in three spikelets positions: apical, central or basal spikelets of the spike. Floret score (dimensionless) was recorded at each sampling for each individual floret developing in each of the three spikelet positions. We only considered for the quantitative analysis of traits determining spike fertility in this paper the floret primordia that reached at least the stage W4.5 (stage when stamen, pistil and carpel primordia are present). For the dynamics of the number of living florets (floret initiation followed by floret death) we only took into account florets that at least reached the stage of W4.5 and a floret was considered dead when it did not show developmental progress (advancement in the floret score) in the following consecutive dissections.

For the purpose of presenting more valuable results we averaged the floret scores of particular floret positions across all the three spikelets (apical, central and basal). While the development F1 in all the three spike positions was very similar (smaller error bars) the distal florets (F2 to Fn) had slower development in apical and basal position compared to that of the central spikelet. So, most of the variation observed due to *Eps-7D* or the temperature and photoperiods were mostly visible in florets F2 and F3.

To determine the overall effects of the *Eps-7D* allele, temperature, photoperiod and their interactions we subjected the data to a full factorial model (a three-way ANOVA) using JMP Pro version 14.0 (SAS Institute Inc., Cary, NC, USA). As the main focus of the paper was to analyse in detail the effect of the *Eps-7D* gene under each of the six growing conditions, we also carried out one-way ANOVA to determine whether the differences between NILs in phenology were significant within each combination of temperature and photoperiod. As the effects of Eps genes are expected to be small, for these analyses we included, in addition to the most conventional levels of probability for significance (i.e. P<0.05; P<0.01; P<0.001) the P-values in each comparison indicating also whenever differences had a P≤0.10 (i.e. significant only at 0.1 probability level) and used P>0.10 and P>>0.1 whenever 0.1>P<0.2 and between 0.21-0.99, respectively.

## Supporting information

Basavaraddi et al., Supplementary figures

## Acknowledgement

Funding for the experimental work was partly provided by projects AGL2015-69595-R, from the Spanish Research Agency (AEI), and IWYP25FP, from the International Wheat Yield Partnership (IWYP). We are grateful to the team of laboratory of crop physiology for assisting with laboratory work. PB held a pre-doctoral research contract from the Agency for Management of University and Research (AGAUR) from the *Generalitat de Catalunya*.

