## Supplementary material for "Wheat developmental traits as affected by the interaction between *Eps-7D* and temperature under contrasting photoperiods with insensitive *Ppd-D1* background": Basavaraddi et al., Supplementary figures

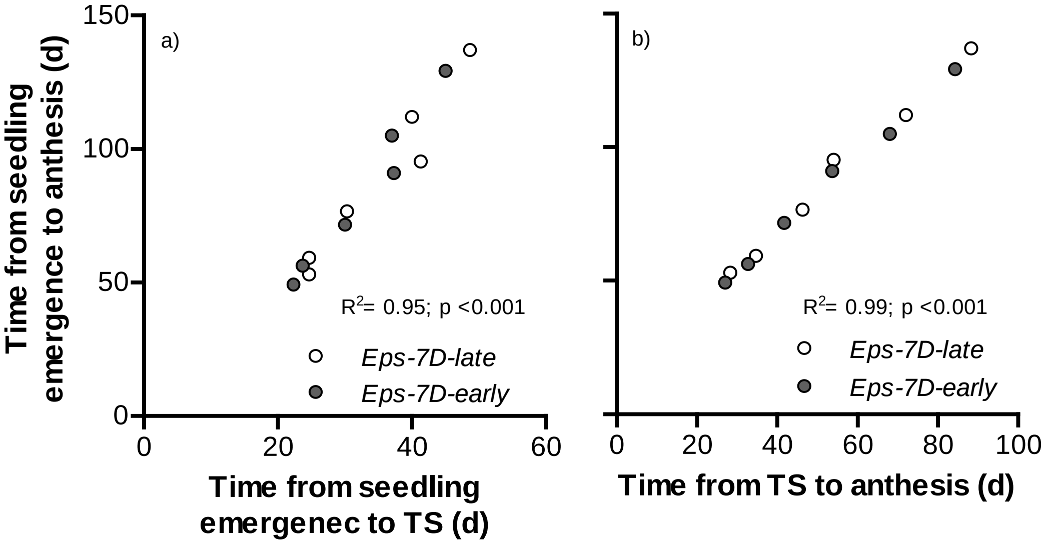


**Supplementary Figure S1.** Relationships between time to anthesis and its component phases: time from seedling emergence to terminal spikelet (TS, a) and time from then to anthesis, i.e. the late reproductive phase (b) for the both the NILs carrying either *Eps-7D-late* or *early* allele under three temperature and two photoperiod regimes.

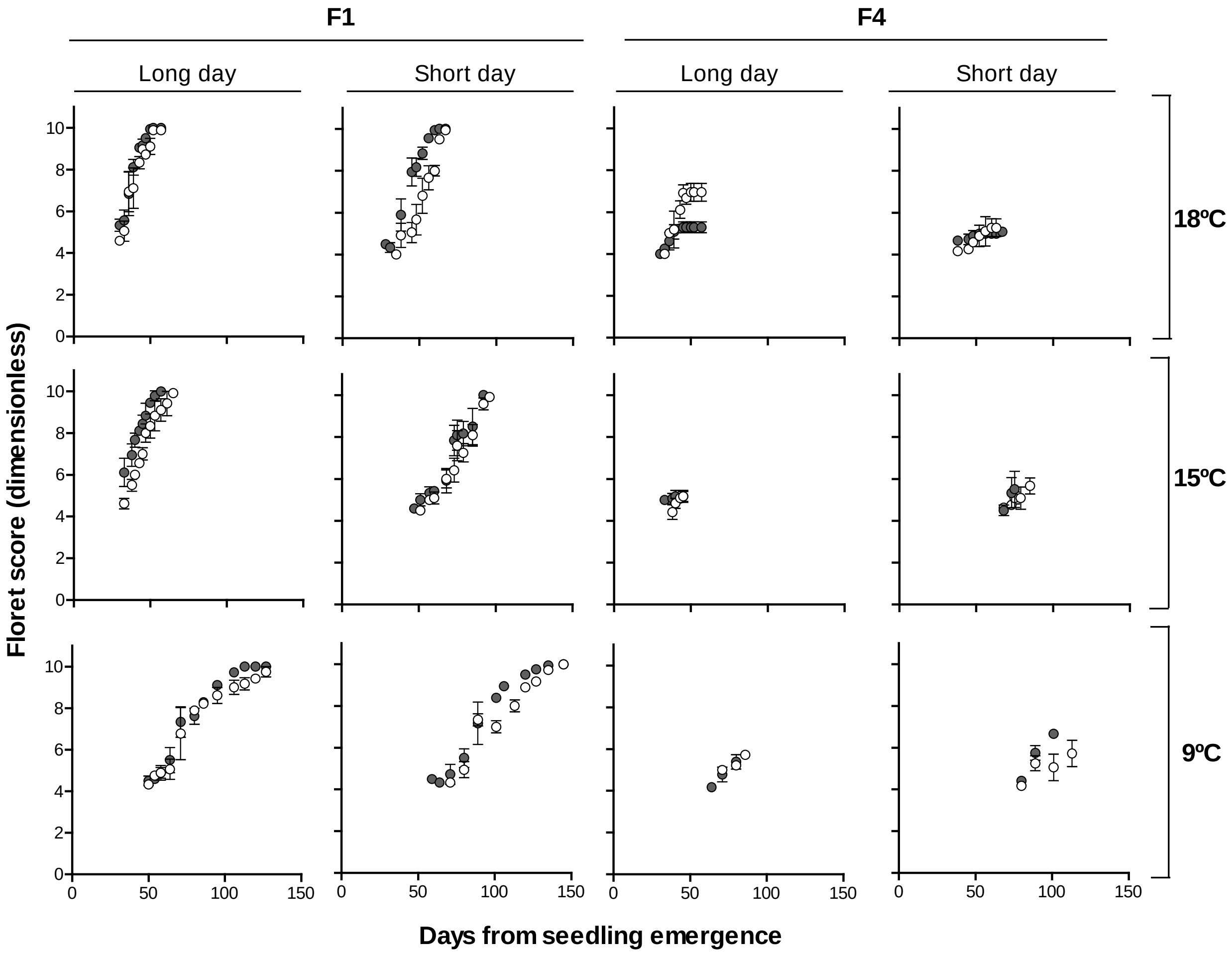


**Supplementary Figure S2.** Relationship between floret development (floret score of the Waddington et al. (1983) scale) and days from seedling emergence for *Eps-7D*-late (open circles) and *early* (closed circles) for floret F1 (left panels) and F4 (right panels) under long and shot day at 18 (upper panels), 15 (middle panels) and 9 ºC (bottom panels). The error bars are SEs of means of floret scores from apical, central and basal spikelets.
